# Teeth Outside the Jaw: Evolution and Development of the Toothed Head Clasper in Chimaeras

**DOI:** 10.1101/2025.04.04.647311

**Authors:** Karly E. Cohen, Michael I. Coates, Gareth J. Fraser

## Abstract

Chimaeras (*Holocephali*) are an understudied group of mostly deep-ocean cartilaginous fishes (*Chondrichthyes*) with unique characteristics that distinguish them from their distant relatives, sharks, skates, and rays. Unlike sharks, chimaeras lack scales and do not have serially replacing rows of serrated teeth crowned with enameloid. Instead, they possess a fused dentition of dentine toothplates. Additionally, male chimaeras develop an articulated cartilaginous facial appendage, the tenaculum, which is covered in an arcade of tooth-like structures. These seeming extraoral teeth remain poorly understood, and their evolutionary origin is unclear. We investigate the development of the tenaculum and its teeth throughout the ontogeny of the Spotted Ratfish, *Hydrolagus colliei*, to assess homology and convergence between this novel craniofacial feature and oral jaws. Our study aims to: (1) describe the development of the tenaculum, (2) assess tenaculum tooth development in comparison to oral teeth and denticles, and (3) characterize the genes and tissues responsible for tenaculum tooth emergence. We found that juvenile male chimaeras develop a full tenaculum before tooth development is complete and that only mature males possess a fully toothed tenaculum. These extraoral teeth emerge from within the tenaculum rather than from the surrounding epithelium. We integrate our developmental data with fossil evidence of the tenacula dentition from the Carboniferous holocephalan *Helodus simplex*. Our findings show that the tenaculum is closely associated with the upper jaw and that tenacula dentition resembles separate shark-like oral tooth whorls more than modified dermal denticles.

**Significance Statement:** The development and evolutionary history of extraoral dentition in vertebrates remain largely unexplored. This study investigates the ontogeny of the male tenaculum, a unique feature of chimaeras, revealing a tooth development pathway similar to the oral dentition in sharks. By integrating fossil data and molecular techniques, we hypothesize that tenaculum teeth are homologous to oral teeth rather than modified skin denticles, providing key insights into the plasticity of odontogenesis and craniofacial diversity in vertebrates.

## Introduction

Living holocephalans are the remnants of a formerly diverse lineage that separated from the ancestry of modern sharks over 385 million years ago (Klug et al., 2023; Coates et al. 2017). As such, holocephalans represent a fundamental division of extant gnathostome diversity and provide unique insights into conditions among early vertebrates. Extant holocephalans are characterized by a distinctive suite of anatomical specializations vastly different from other chondrichthyans (Didier et al., 2012; Didier et al., 1998; Ferrando et al., 2016; Inoue et al., 2010; Johanson et al., 2021). They mostly lack denticles and do not have a replacing arcade of separated teeth; instead, they have a tooth-plate composed of only dentine. Among their distinctive features, one standout is the tenaculum, a club-shaped, articulated, cartilaginous facial appendage studded with an arcade of tooth-like structures found only in male chimaeras (Didier et al., 2012; Didier et al., 1998). The tooth-like structures are exposed and visible when the tenaculum is extended (raised out of the head); and when not in use, the unit sits in a central recess behind the eyes (Figure 1). The tenaculum is an unmistakable structure on the front of the male head, it is typically not pigmented compared to the surrounding head color pattern. The function of this sexually dimorphic structure is unknown, although video records of rare mating behaviors suggest the toothed tenaculum is used by males to grip female pectoral fins during copulation. Some of the first descriptions of this dimorphic behavior was noted by Dean (1906) who described scars present on the dorsal surface of female *Hydrolagus colliei* caused by the scraping of male tenacula teeth during courtship.

**Figure 1:**
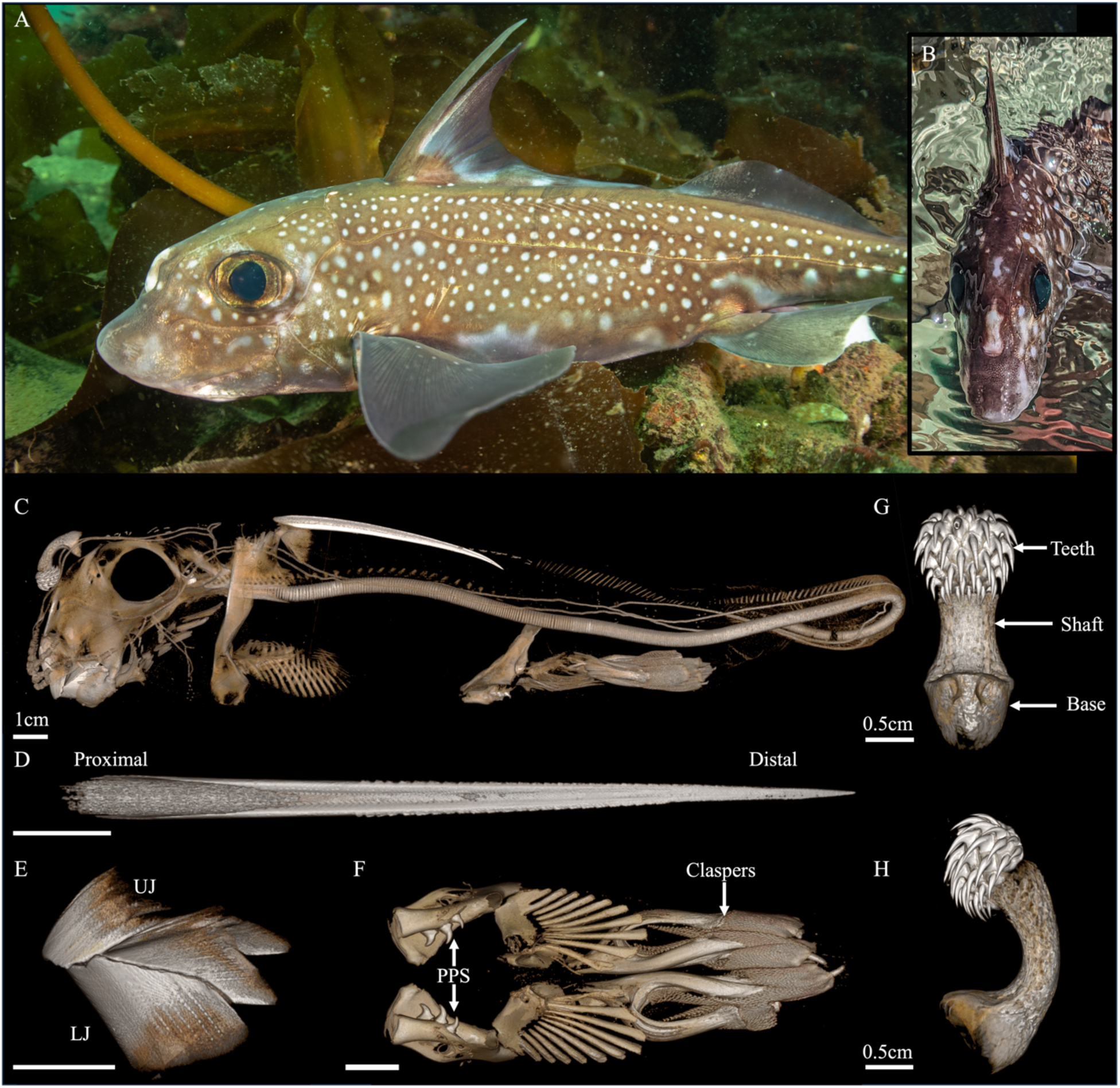
Basic anatomy of *Hydrolagus colliei*. (A,B) Photograph of adult, male *H. colliei* in Puget Sound. (Photograph used with permission from Tiare Boyes) (C) Micro-CT scan of male *H. colliei*. (D) Micro-CT scan volume of dorsal spine in ventral view, highlighting serrations along the spine’s length. (E) Volume rendering of modified beak, hypermineralized regions visible as beaded columns. (F) Volume rendering of claspers and pre-pelvic anatomy. The pelvis is studded with six large denticles, and the claspers are covered in hundreds of small, rhomboid-shaped denticles. (G, H) Segmentation of the adult tenaculum, with teeth colored to highlight arrangement. Scale = 1 cm.

A number of anatomical specializations in male chimaeras are directly tied to courtship and mating (Didier et al., 2012, Figure 1). Males possess multiple grasping structures, including a pair of pelvic claspers covered in sharp denticles, a pre-pelvic tenaculum positioned just anterior to the pelvic fins, in addition to the cephalic tenaculum (Leigh-Sharpe 1922; Patterson 1965; Didier 1995; Didier et al., 2012; Figure 1). These highly specialized and diverse structures are found across all extant holocephalans, underscoring the evolutionary investment in reinforced mating behaviors within the extended lineage. Beyond their reproductive morphology, chimaeras diverge from other chondrichthyans in possessing a beak-like arrangement of hypermineralized tooth plates rather than individual, replaceable teeth. This feeding apparatus consists of three paired plates—mandibular, vomerine, and palatine—each exhibiting distinct mineralized tissues, including trabecular dentine and hypermineralized tritors, which provide the durability needed for durophagy (Johanson et al., 2020; 2021; Smith et al., 2019). Despite these departures from other cartilaginous fishes, holocephalans have retained at least one deeply conserved feature: the dorsal fin spine. Present in all living species and often venomous, the rigid spine is a feature retained from the earliest members of the chondrichthyan clade, extending back for at least 436 million years (Zhu et al., 2022).

Fossil holocephalans exhibit diversity absent in modern lineages. For instance, some individuals possess oral dentitions with serially organized molariform teeth (Stahl 1999). It has long been suggested that the transformation and development of these individual tooth units eventually led to the tooth plate dentition of modern lineages (Finarelli & Coates, 2011; Johanson et al., 2021; Patterson, 1965). In addition, extinct chimaeras featured various extraoral structures adorned with teeth, including a median frontal clasper (Patterson, 1965; Stahl 1999). Patterson (1965) suggested that toothed frontal claspers (tenacula) were originally present in both sexes, based on fossil evidence of early Chimaeriformes. Therefore, it was postulated a defensive function may have preceded the restriction to males for a copulatory function (Patterson, 1965): an intriguing notion although unsupported thus far by the proliferation of Palaeozoic forms (Stahl 1999) evident at fossil localities such as Bear Gulch (Grogan, 1993; Grogan et al., 2012).

Tenacular teeth are presumably dermally derived, similar to the dermal denticles in other cartilaginous fishes. Yet, without early developmental or genetic data it remains unclear if the evolutionary origin of tenacula teeth lies in the evolution of oral teeth or if the tenaculum represents convergent modification of body denticles. The broader debate surrounding odontodes—the mineralized structures composed of dentine found both externally and internally in vertebrates—adds complexity to this question. While teeth are often considered a specialized subset of odontodes, recent perspectives challenge the assumption that teeth evolved directly from dermal denticles, as early vertebrate scales differ substantially from those of modern sharks (Berio & Debiais-Thibaud, 2021; Fraser et al., 2009; Haridy et al., 2019; Nicklin et al., 2024; Smith & Coates, 2000). Instead, early vertebrate skeletal tissues exhibit a mosaic of structures that complicates straightforward homology between teeth and scales. Chondrichthyans (shark, skates, rays, and chimaeras) have long been studied for their dental characteristics as they provide ample opportunity to investigate the evolutionary or developmental origin of teeth (Fraser et al., 2009; Fraser & Hulsey, 2020; Hulsey et al., 2020; Rasch et al., 2016; Tucker & Fraser, 2014; Brownstein et al. 2024). If tenacular teeth are homologous to oral teeth, they may provide a missing link in the evolution of mineralized dental structures beyond the oral cavity. Conversely, if they are instead modified body denticles, their independent evolution could highlight the plasticity of dermal odontogenic mechanisms in chondrichthyans.

In this study, we provide the first account of complete tenaculum development and ontogeny in chimaeras. We take advantage of an uncommon set of ontogenetic stages from the Spotted Ratfish, *Hydrolagus colliei*; one of approximately 50 species of extant Chimaera, across 3 families (Brownstein et al., 2024; Didier et al. 2012; Stahl 1999). Using histology and μCT scanning we assess how the tenaculum and its associated tooth set develops. We ask whether tenacula tooth development is more similar to that of the oral dentition or follows a pattern of development similar to the skin denticles seen in other elasmobranch lineages. Through developmental stages of tenaculum ontogeny and immunohistochemistry we characterize the molecular signature of tenaculum tooth emergence and renewal, aiming to establish connections between modern and extinct dental features.

## Results

### Tenaculum development

Tenaculum development initiates during embryonic development (*in ovo*) and appears to be a default morphological structure observed in both embryonic males (Figure 2A&B) as well as females (Figure 2 C&D). In males, the tenaculum continues to grow through latter stages of juvenile development; from a small cellular condensation in embryos to a pimple-like structure situated at the midline between the orbits and the rostrum (Figure 2A; Figure 3A&B). As the tenaculum continues to grow, the characteristic rod gradually takes shape, elongating into a pill form (Figure 3 C&D). CT scans revealed that, at this stage, there are only minimal levels of mineralization and true cartilage surrounding the outer surface of the tenacula rod (Figure 3 C&D). By way of comparison, we examined the earliest stages of development in *Callorhinchus milii* (the ‘elephant shark’ chimaera, Venkatesh et al. 2014) and observed that the tenaculum arises in a manner broadly analogous to that seen in *H. colliei.* In both CT scans and histology, we find the callorhinchid tenaculum starts as a small epithelium thickening just behind the eyes. Micro-CT scans and histological analysis reveal that the callorhinchid tenaculum originates as a modest epithelial thickening situated just posterior to the orbital region. This initial condensation enlarges and subsequently begins to mineralize, assuming the appearance of a pimple-like protrusion (see Supplementary Figure S1). Notably, the base or proximal end of the rod exhibits a more bulbous shape compared to the anterior or distal end. Furthermore, we observe the insertion of muscle fibers at the proximal end from both sides. Although the exact origin of these muscle fibers remains unclear, contrast CT imaging suggests recruitment from the adductor series.

**Figure 2:**
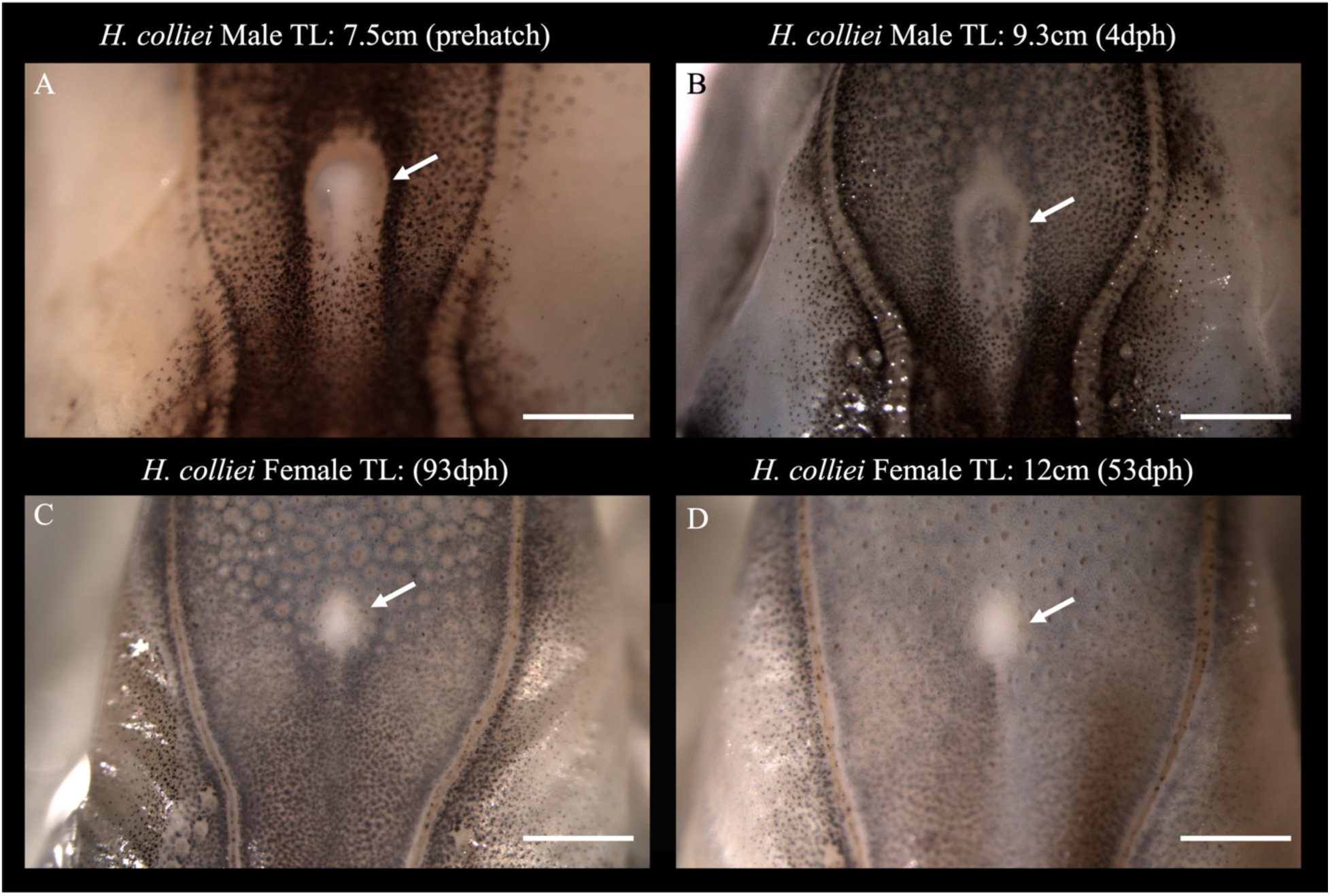
The rudimentary tenaculum (pimple) is a morphological default present in both embryonic male and female *H. colliei*, retained later only in males. Condensations of the rudimentary tenaculum are apparent as white thickenings (later cartilage, white arrow) in the superficial mid-interorbital region of embryonic Spotted Ratfish. (A, B) In males, this condensation becomes more pronounced and furrowed at the apical tip. (C, D) In embryonic females, the condensation is less pronounced and appears as a white spot in the region of the male tenacula tip.

**Figure 3:**
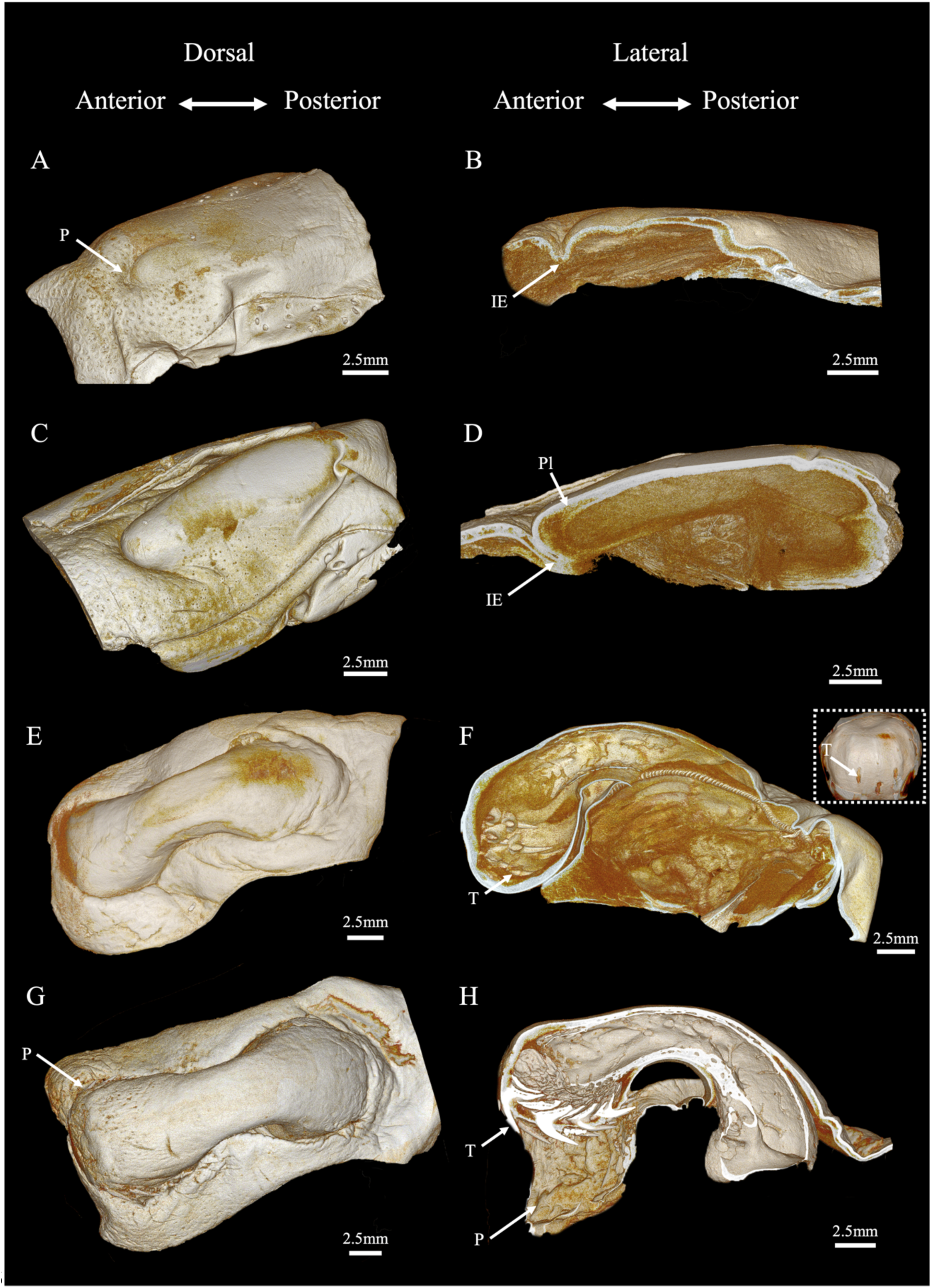
Development of the tenaculum in *H. colliei*. (A–G) CT dorsal view of developing tenaculum from 20–85 cm SL. (B–H) Lateral section revealing internal anatomy. IE = invaginating epithelium; P = pocket; T = teeth.

At approximately 25 cm total body length (TL), the tenaculum undergoes elongation and widening, eventually erupting through the surface skin on the ratfish’s forehead (Figure 3 C&D). As the tenaculum widens, a central pocket forms around the distal end, which continues to accommodate the tenaculum through juvenile development. Through the whole of development, the front of the tenaculum is tightly adhered to the epithelial pocket and cannot be extended or removed. During this stage, we observe the differentiation and separation of epithelial tissue, creating a distinct line of tissue throughout the anterior bulb.

Tenaculum tooth development first becomes evident in specimens ranging from 30 to 50 cm total length (TL, Figure 3 E-F). Teeth emerge in sequential rows, originating from lamina tissue formed during earlier ontogenetic stages. On average, seven rows develop, with the most proximal rows forming first and exhibiting more advanced tooth stages, while earlier-stage teeth appear in more distal rows. The base of each tooth overlaps with its adjacent neighbor however, the tooth crowns do not overlap but rather interdigitate across rows. They are curved, conical shape, retain a central pulp cavity, and highly mineralized tips.

Teeth gradually erupt through the closed cartilaginous anterior bulb of the tenaculum (Figure 3G–H). The first two teeth appear at the midline of the initial row, followed by the sequential emergence of additional teeth. This process significantly widens the epithelial pocket on the surface. As the teeth protrude, the tenaculum bulges outward. The tenaculum is now free from the surrounding epithelium and can either be erected or retracted. Once all teeth have erupted, individual recesses become visible in the pocket, allowing the teeth to slot into place when the tenaculum retracts. The cartilage surrounding the base of each tooth expands into a teardrop shape (Figure 3 G&H).

Histological examination of juvenile through adult stages reveals that the tenaculum has a core of dense mesenchymal tissue surrounded by a layer of mineralized cartilage (Figure 4). In early juveniles, the cells proliferate and organize to form a small cartilaginous nub (Figure 4 A&B). This nub, when viewed from the surface, presents itself as a mineralized white spot or “pimple” on the fish’s forehead. Histological sections indicate that the outer epithelium covering is sequestered and folded underneath the developing tenaculum (Figure 4 A&B; IE). The developing tenaculum has a central core primarily composed of undifferentiated mesenchymal cell types (Figure 4 D&E; M). Eventually, strings of epithelial condensations are found inside the anterior end of the tenaculum representative of a dental lamina (Figure 4 E; DL). As the tenaculum continues to grow the inner core regulates into hyaline cartilage with a layer of mineralized tissue surrounding the entire rod (Figure 4 G: MC). The tooth-bearing anterior end of the tenaculum is less mineralized than the rest of the structure, with teeth extending from inside to outside, embedded in a dense matrix of mesenchyme (connective tissues) and cartilage (Figure 4G). Protein expression analyses (immunohistochemistry) of early and late stage developing tenacula reveal expression of SOX2 and B-CAT in these streams of epithelial tissue. These signals continue to progress throughout the differentiation of the tenaculum and early stages of tooth morphogenesis (Figure 4 C,F,I). This signal is observed in the earliest stages of tenaculum growth and development with the first signs of cellular condensation occurring on the cranium. Prior to tenaculum development we do not see such expression patterns.

**Figure 4:**
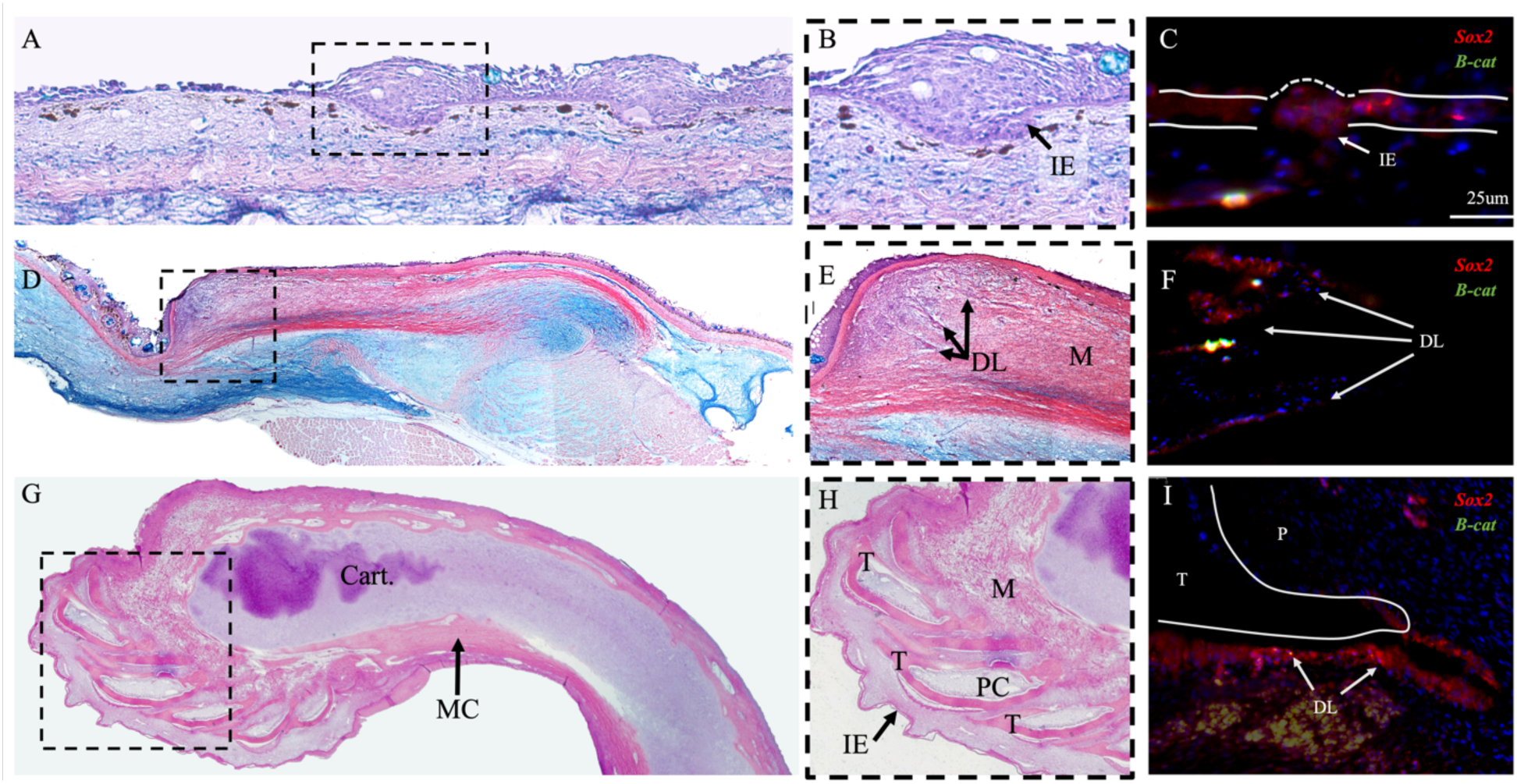
Histology and gene expression of developing tenaculum. (A, B) Cellular condensation marks the first signs of tenacular development as epithelial layers incorporate. (C) Tissue staining reveals gene expression patterns of SOX2 and B-CAT in early developing tenacula. (D, E) The tenaculum assumes its rod-like shape, and a dental lamina (DL) forms. (F) Gene expression analysis indicates odontogenic potential in the tenaculum. (G, H) Adult tenaculum structure, showing serially arranged teeth embedded in mesenchyme and cartilage. IE: internal epithelum, DL: dental lamina, Cart: cartilage, T: tooth, PC: pulp cavity, M: mesenchymal cells

The adult tenaculum is a club-like structure composed of a dense cartilaginous core encased in a fibrous connective tissue sheath. The distal surface is embedded with numerous small, recurved denticles arranged in about seven rows. The teeth retain their pulp cavity even after eruption (Figure 4G & 5A), and the outer surface of the tooth-bearing area is covered by a thick layer of sponge-like epithelial tissues devoid of goblet cells or tooth buds, allowing the teeth to flex slightly (Figure 5A). tooth Following each tooth into the tenaculum is a string of epithelial tissue that wraps around the base of each tooth (Figure 5 B&C). Adult tenacula vary in the total number of teeth and rows, though the average tooth size within each row remains consistent. Teeth at the center of each row are larger than those at the lateral edges (Figure 5D &E). Additionally, smaller, unaligned teeth are present on the ventral surface of the anterior bulb.

**Figure 5:**
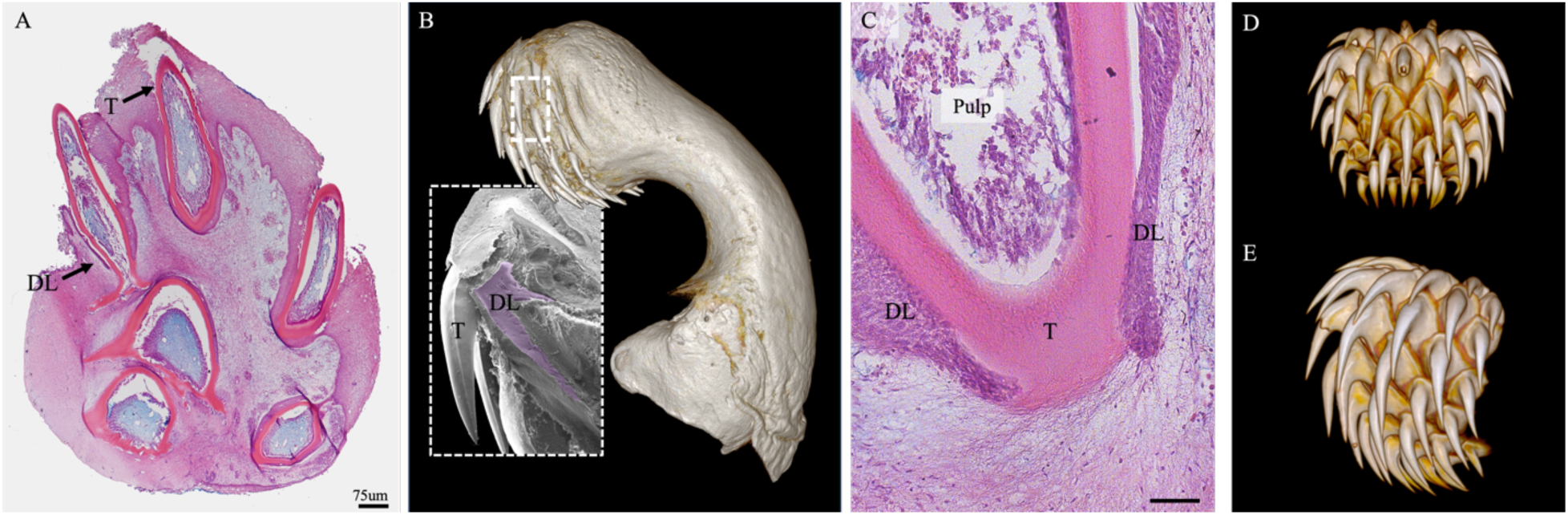
Adult tenaculum morphology. (A) CT scan and scanning electron micrograph (inset) of adult morphology, showing the positioning of the dental lamina. (B) Cross-section of tenaculum, with exposed teeth retaining pulp cavities. (C) Histological section showing the dental lamina surrounding a tenaculum tooth. (D) Frontal view of adult tenaculum and (E) lateral view showing the relative size and overall of adult teeth.

### Tenaculum growth vs. clasper growth

We anticipated parallel growth between the tenaculum and body length; however, our investigation revealed a distinct relationship between the tenaculum’s size and the development of the pelvic claspers. Smaller males with fully developed pelvic claspers exhibited significantly larger tenacula than some of the longer males with less developed claspers. Over ontogeny, claspers develop numerous small diamond-shaped denticles. Histology revealed that these denticles are mineralized with dentin and an empty pulp cavity. Notably, no replacement denticles were observed beneath the surface of the odontode-bearing epithelium (see Supplementary Figure S2). In the earlier stages of clasper denticle development, we observed small placodes condensing and emerging from the skin as tissues differentiated and underwent mineralization. Unlike tenaculum teeth, there is no lamina-like structure associated with clasper denticles.

## Discussion

The process of tooth development and emergence is highly conserved across gnathostomes (jawed vertebrates), but the chimaeras present the first example of a dental lamina outside of the jaw and offer new insights into the possibilities of gnathostome dental diversity. The dental lamina begins as an invagination of oral epithelium and its early establishment in gnathostome jaws is crucial for proper tooth emergence and regeneration (Fraser et al., 2009, 2019; Huysseune, 2006; Smith et al., 2009). In chondrichthyans, continuous dental replacement and the alternating pattern of tooth emergence is controlled by genetic expression through this ectodermally derived string of tissue (Cooper et al., 2022; Fraser et al., 2019; Nicklin et al., 2024; Rasch et al., 2016; Schnetz et al., 2016). Sharks and their relatives (elasmobranchs) are covered in tooth-like scales known as dermal denticles, but these do not emerge or replace by means of a dental lamina (Reif 1980, 1982). Rather, concentrated expression of an odontogenic network is responsible for sequential generations of denticle replacement. Unlike elasmobranchs, modern chimaeras lack extensive scale-cover (Patterson 1965; Didier 1995; Stahl 1999). Remaining denticles are limited to the pre-and-post pelvic claspers, the dorsal spine in some very large males, and those ephemerally present on the cranium in hatchlings (separate to the tooth-like whorls present on the head tenaculum; Didier et al. 2012; Finarelli & Coates 2014).

While teeth and denticles in elasmobranchs share several key genetic and physical traits, teeth uniquely develop from a dental lamina in the oral epithelium. The present study shows that the tenaculum of *Hydrolagus colliei* develops from an island of cells that could potentially be oral epithelial cells found on its forehead (further work will elucidate the cellular lineage origins of these cells), palaeontological evidence has provided context for this hypothesis (see below). Expression of Sox2, B-Cat, and other key genetic markers are comparatively stronger in this thickened forehead epithelium, likely conferring odontogenic potential to the underlying dental lamina-like tissue. Teeth continue to develop along and in response to this string of epithelial tissue throughout development. Furthermore, in adults, this tissue dives around each of the tenacula teeth in a similar form to what is observed in the dentitions of sharks and batoids. The presence of this dental lamina in the tenaculum of *H. colliei* challenges traditional views on the spatial segregation of teeth from denticles (Cooper et al., 2018, 2022; Nicklin et al., 2024). We suggest that the origin of this diversification lies in extinct chimaeroid fishes where the tenaculum first evolved and adds substantial evidence to previous discussion about the evolutionary origin of holocephalan tenaculae first offered on the basis of morphological data alone (Coates et al. 2021).

The cranial skeleton of the Pennsylvanian (Moscovian; around 315 million years old) holocephalan *Helodus simplex* bears the oldest and most primitive known example of a tenaculum (Coates et al., 2021; Fig. 6A-C). The tenacular cartilage (the stem or rod) lies above the ethmoid region, which itself appears to represent an intermediate stage in the evolution of the highly specialized preorbital region of modern chimaera crania (de Beer, 1937; Patterson, 1965; Didier et al., 2012; Didier, 1995; Fig. 6D). The evolutionary origin of the tenacular rod is unclear, other than its proximity to the ethmoid roof of which it might be considered a subdivision or outgrowth. However, the denticles of the *Helodus* tenaculum are remarkably tooth-like (Fig. 6C). These teeth are arranged in a tightly packed whorl that coils through the front of the tenacular cartilage. Furthermore, these tenacular teeth have roots that resemble those of the mandibular teeth, although the bicuspid crowns are markedly different from the pillow-shapes of the palatal and mandibular dentition (Fig. 6B). Here, it is worth mentioning that *Helodus* mandibular teeth exhibit pre-conditions for the dental plates of modern chimaeras (Didier, 1995; Johanson et al., 2021; Stahl, 1999), with separate teeth (cf. elasmobranch tooth sets) consolidated into massive batteries occupying each quarter of the gape (Coates et al., 2021). The *Helodus* tenaculum extends over the entire length of the ethmoid region, and the apical tooth whorl (which is rather small compared to neighbouring teeth) sits within a symphysial gap between the left and right upper jaw tooth batteries (Fig. 6A; Coates et al., 2021). This extraordinary tenacular length, relative to modern examples, is not unique and might, in fact, represent a primitive condition: similarly elongate tenaculae are known in Mesozoic holocephalans (Patterson, 1965) including genera such as *Squaloraja*, *Acanthorhina*, and *Metopacanthus* (Stahl, 1999). These, too, bear a wide variety of denticles, but detailed descriptions have not been completed.

**Figure 6:**
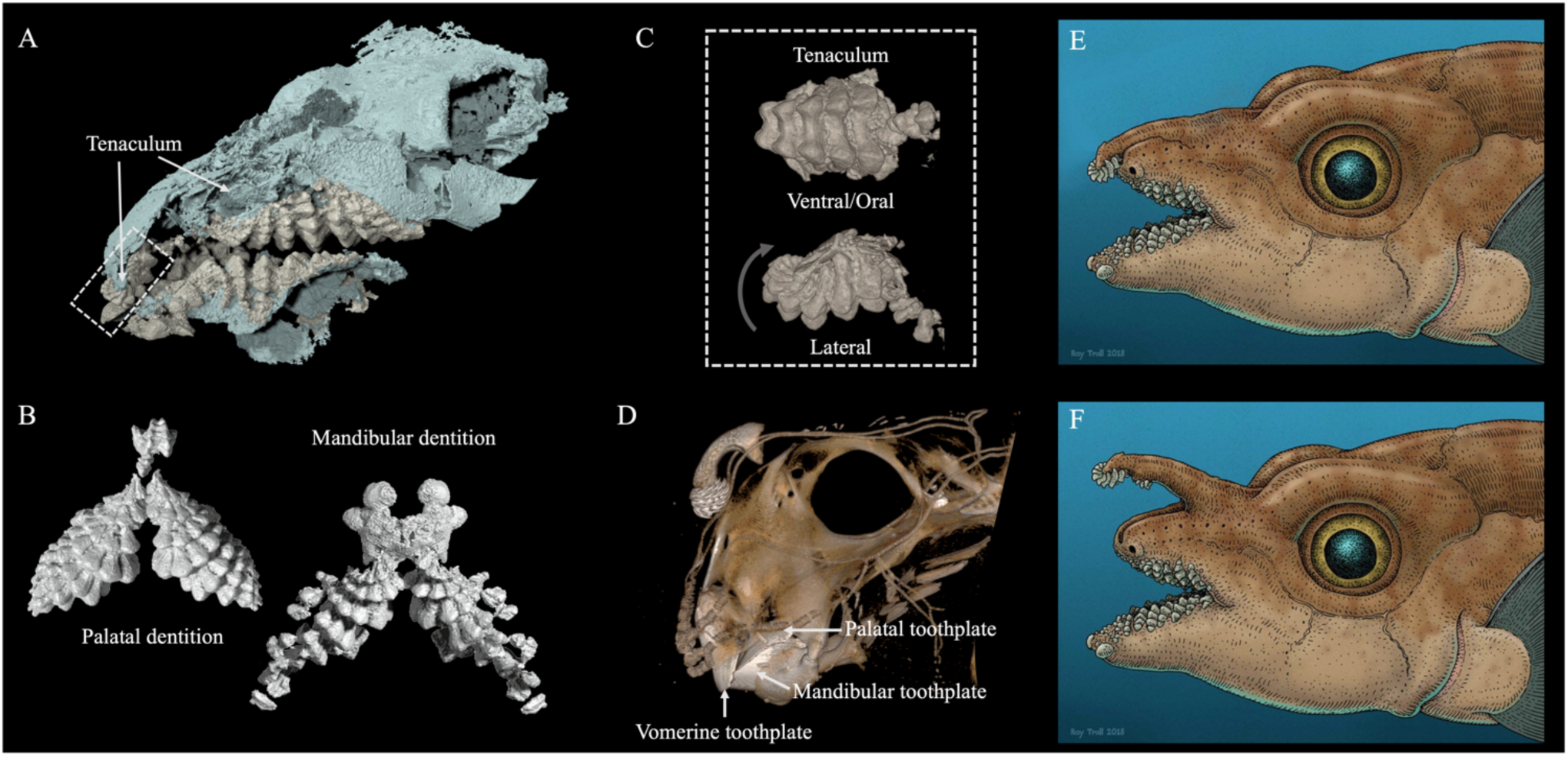
Origin of the modern tenaculum. A) The Carboniferous stem-holocephalan *Helodus simplex* (see Coates et al. 2021 for details) cranium, partly crushed, in anterolateral view showing elongate tenaculum with tenacular teeth positioned anterior to palatal dentition and (when depressed) seated in gap between anterior mandibular teeth. B) *H. simplex* tenacular tooth whorls (large and small). C) *H. simplex* dentition, palatal and mandibular, showing closely interdigitated tooth families with lingual to labial serial replacement akin to dental arcades present in modern sharks and rays. D) *Hydrolagus colliei*, the spotted ratfish, showing the short tenaculum of modern chimaeroids and tooth plate dentition. The tenaculum retains ancestral features such as individual tooth units, reflecting its origin in the oral jaw dentition rather than as a modification of dermal denticles. E & F) Reconstruction of *H. simplex* (used with permission from Ray Troll) showing likely elevated and depressed positions of elongate primitive tenaculum.

Here we suggest that in Palaeozoic taxa such as *Helodus*, the close proximity of the tenacula tip with the upper jaw dentition allowed for an epithelial connection, at least during development, promoting a transfer of dental competence onto the tenacula appendage. This, in turn, seems likely to be a legacy of these teeth originating, evolutionarily, from the front of the gape. Further, we argue that this retention and/or transfer, derived within the stem lineage of today’s Chimaeriformes, led to the development of what we now see as oral teeth outside of the jaws in extant holocephalans. Subsequently, under differing selection regimes, the tenacular teeth and jaw teeth became increasingly dissociated and adopted new and separate functions. A curious result is that cyclical tooth replacement, widely considered to be a defining characteristic of chondrichthyans, in chimaeras is retained only in the tenaculae while distinctly absent in the mandibular dentition (Didier, 1995; Johanson et al., 2021; Stahl, 1999).

The tenaculae of extant chimaeras are morphologically diverse, and this probably reflects their as yet undefined roles in reproduction. In species like *Harriotta raleighana*, tenacula teeth are long and slender, whereas in *Callorhinchus milli,* they are short and squat (Carrier et al., 2004; Didier et al., 2012; Didier et al., 1998; Nakayama et al., 2020). Notably, some chimaeras exhibit occluding tenacula teeth located outside, within the pocket housing the tenaculum (i.e., *Callorhinchus milii*), in stark contrast to *H. colliei* where teeth are confined to only the tenacular bulb. This pattern indicates that odontogenic cells capable of tooth formation are distributed even further across the forehead and are not restricted to tenacular territory. Despite differences in tooth morphology, all tenacula share the defining characteristic of a serially replacing dentition, a capability (as noted above) lost in the oral jaws of modern chimaeras. While a dental lamina is not always essential for the initial emergence of teeth (Vandenplas et al., 2014), and several vertebrates generate odontic structures without one, it is required for sustaining a continuous dentition (Fraser et al., 2020).

Chimaera dermal denticles are structurally and histologically distinct from tenacular teeth. Dermal denticles surrounding the dorsal spine lack a mineralized crown and consist solely of a base, while pelvic clasper denticles, like those in elasmobranchs, are embedded in the skin without lamina-like tissue. In contrast, tenacular teeth mineralize in a manner like true teeth in other chondrichthyans, with the crown forming before the base. These teeth retain a pulp cavity, contain multiple layers of dentin, and may be capped with enameloid. The presence of dentin and pulp, which are absent in dermal denticles, suggests a structural and functional distinction. The tenaculum is considered a sexually dimorphic structure, but its development appears to be at least partially hormonally regulated. While males develop a fully formed tenaculum with functional teeth, females may form a rudimentary shaft that never develops teeth or resembles the adult male structure (Leydig 1851, Dean 1906). This pattern suggests that the tenaculum follows a default developmental pathway that matures only in males while regressing in females (Figure 2). A similar trend is seen in other sexually dimorphic dermal structures in chimaeras, where denticle appearance is often linked to reproductive traits or body size.

In summary, our findings challenge the idea that the tooth-like denticles of the tenaculum are modified dermal denticles. Rather, we argue that tenacular denticles are representations of true teeth outside of the jaw whose origin lies in facial epithelium that was once competent for the establishment of the oral dentition. Coupled with recently described material from fossils and new phylogenies, these data shed light on the evolution of an extraordinary morphological innovation: a rare example of the kind of evolutionary developmental tinkering long theorized about (Lieberman & Hall, 2006) but rarely substantiated with experimental and comparative data.

## Materials and Methods

### Specimen collection

Samples of adult and juvenile *Hydrolagus colliei* were collected during trawls in the San Juan Channel, Friday Harbor Washington, USA. These trawls were deployed from April 2022 - August of 2023. All samples collected with permission under a University of Washington animal care and use protocol (IACUC 4238-03).

### Tenaculum and body ontogenetic morphology

We used micro-computed tomography (µ-CT) to examine the morphology and development of the tenaculum in *H. collei*, scanning both unstained and dye-enhanced specimens. In total, we analyzed 40 specimens ranging from 25 to 80 cm total length (TL). For whole-body morphometrics, we µ-CT scanned 16 specimens without contrast. These scans were conducted at the University of Florida using a Nano-CT GE V|TOME|X M 240 at 28–25 µm resolution, reconstructed in VG Studio, and exported as .tiff files for segmentation in 3D Slicer. To visualize non-mineralized tissues and developing tooth buds, we dissected 12 additional tenacula for dye-enhanced CT scanning. These were immersed in a 3% phosphotungstic acid (PTA) and ethanol solution for one week with continuous agitation to improve dye penetration. Scans were performed at the Friday Harbor Laboratories Karel F. Liem Bio-Imaging Center using a Bruker Skyscan 1173 with a voxel size of 6.9–12.1 μm, a voltage of 55 kV, an amperage of 133 μA, and an exposure time of 1.175–1.350 s. We processed all scans using 3D Slicer and the SlicerMorph toolkit for segmentation, visualization, and morphological measurements (Kikinis et al., 2014; Rolfe et al., 2021).

### Histology

Adult and juvenile samples (n=7) were decalcified using either 0.5 M EDTA in water for 1–2 weeks until softened or Cal-Ex for 12 hours at 4°C (for adult tissues). Following decalcification, samples were dehydrated in ethanol, cleared with xylene, and embedded in paraffin. We sectioned sagittal paraffin-embedded samples (6–7 μm thick) using a Leica RM2145 microtome. Slides were stained with hematoxylin and eosin, mounted with DPX (Sigma), and imaged using a BX51 Olympus compound microscope equipped with an Olympus DP71 camera. Additional slides were prepared for immunohistochemistry and stained for SOX2, PCNA, and activated β-catenin. These sections were mounted with Fluoromount, sealed with fluoromount, and imaged with BX51 Olympus compound microscope fitted with an OlympusDP71 camera.

### Statistical analysis

We measured tenaculum length, tooth count, individual tooth length, volume, and surface area for all adult and subadult specimens. Total length and clasper length were also recorded, with the former measured in ImageJ from DSLR images (Canon Mark III, 180mm macro lens) and the latter assessed in 3D Slicer. To evaluate tenaculum size variation among adults, we conducted a two-way ANOVA with clasper development and total length as independent variables. The analysis met assumptions of independence, homogeneity of variances, and normality of residuals. The significance threshold was set at α = 0.05. Data and analysis code are available upon request.

## Acknowledgements

We would like to thank Jakobb Bueche and Eric Loss, Marine Operations Supervisor and Captains of the *R/V Kitiiwake* for their endless support in deploying trawls. Kristy Kull for her fishing expertise. Thank you to Emily Carr for initial probes into the morphology of the Ratfish during an NSF REU 2020. We thank Fidji Berio and Plant Ocean aquarium, Montpellier, France, for the gift of the embryonic *H. colliei* specimens and Emma Bernard of the Natural History Museum, UK, for access to H. simplex specimens. Additional thank you to Friday Harbor Laboratories for the New Faculty Fellowship to GJF, the Stephen and Ruth Wainwright Endowment to KEC, and National Science Foundation Awards IOS: 2128032 to GJF; EAR: 2218892 to MIC.

---

**Figure S1.**
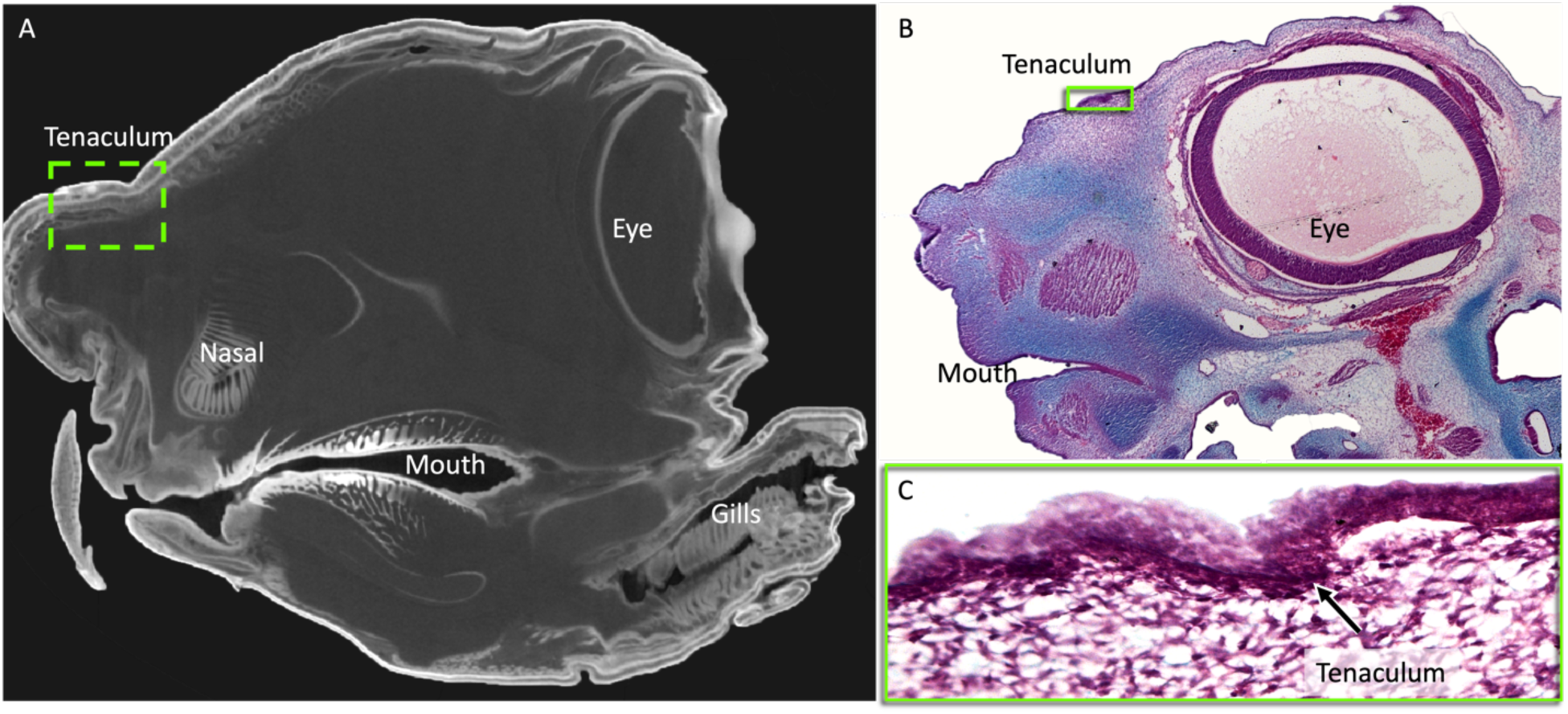
Early development of the tenaculum in *Callorhinchus milii*. (A) Micro-CT scan of a late-stage embryonic *C. milii* head showing the position of the developing tenaculum (green box), located anterior to the eye and dorsal to the nasal capsule. (B) Histological section through a comparable plane illustrating the overall cranial anatomy, with the early tenaculum region boxed in green. (C) Higher magnification of the boxed region in (B), highlighting the tenaculum as a dense cellular condensation (black arrow), indicating the initial stages of its formation. Panels A–C are oriented similarly, and the green boxes in each panel correspond to the location of the developing tenaculum.

**Figure S2.**
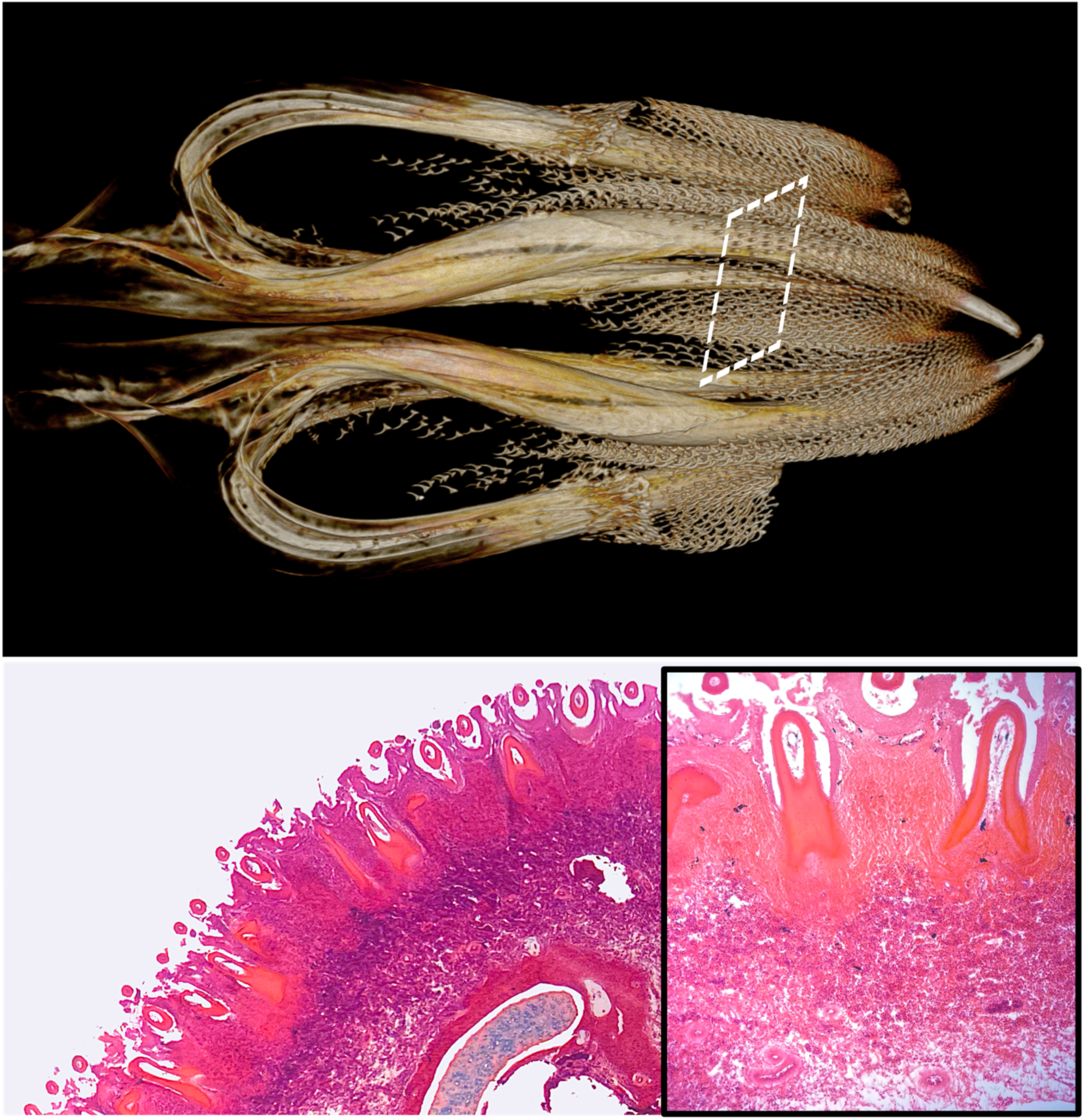
Morphology and histological organization of the denticle-covered claspers in male *Hydrolagus colliei*. Top: CT scan of male claspers showing dense coverage of denticles (white dashed box indicates region shown below). Bottom left: Histological section through the clasper shaft illustrating the organization denticles within the epidermis. Bottom right: Higher magnification of the boxed region, highlighting the mature denticle morphology and lack of replacement denticles. These structures are histologically distinct from the developing tentacular teeth.

## References

1. Klug C, Coates M, Frey L, Greif M, Jobbins M, Pohle A, Lagnaoui A, Haouz WB, Ginter M (2023) Broad snouted cladoselachian with sensory specialization at the base of modern chondrichthyans. Swiss J Palaeontol 142:2. 10.1186/s13358-023-00266-6

2. Coates MI, Gess RW, Finarelli, JA, Criswell KE, Tietjen K. (2017) A symmoriiform chondrichthyan braincase and the origin of chimaeroid fishes. Nature, 541(7636), 208–211. 10.1038/nature20806

3. Didier DA, Kemper J, Ebert D (2012) Phylogeny, biology, and classification of extant holocephalans. In Carrier JC, Musick JA, Heithaus MR, eds. Biology of Sharks and Their Relatives, Second Edition, pp. 97–122 (CRC Press, Boca Raton, FL). 10.1201/b11867-6

4. Didier DA, LeClair EE, Vanbuskirk DR (1998) Embryonic staging and external features of development of the chimaeroid fish, *Callorhinchus milii* (Holocephali, Callorhinchidae). J Morphol 236:25–47. 10.1002/(SICI)1097-4687(199804)236:1<25::AID-JMOR2>3.0.CO;2-N

5. Ferrando S, Gallus L, Gambardella C, Croce D, Damiano G, Mazzarino C, Vacchi M (2016) First description of a palatal organ in *Chimaera monstrosa* (Chondrichthyes, Holocephali). Anat Rec 299:118–131. 10.1002/ar.23280

6. Inoue JG, Miya M, Lam K, Tay BH, Danks JA, Bell J, Walker TI, Venkatesh B (2010) Evolutionary origin and phylogeny of the modern holocephalans (Chondrichthyes: Chimaeriformes): A mitogenomic perspective. Mol Biol Evol 27:2576–2586. 10.1093/molbev/msq147

7. Johanson Z, Manzanares E, Underwood C, Clark B, Fernandez V, Smith M (2021) Ontogenetic development of the holocephalan dentition: Morphological transitions of dentine in the absence of teeth. J Anat 239:704–719. 10.1111/joa.13445

8. Dean B (1906) Chimaeroid Fishes and Their Development. (Carnegie Institution of Washington, Washington, DC).

9. Leigh-Sharpe WH (1922) The comparative morphology of the secondary sexual characters of elasmobranch fishes, memoirs IV and V. J. Morphol. 36:199–243.

10. Patterson C (1965) The phylogeny of the chimaeroids. Philos Trans R Soc Lond B Biol Sci 249:101–219.

11. Didier DA (1995) Phylogenetic systematics of extant chimaeroid fishes (Holocephali, Chimaeroidei). Am. Mus. Nat. Hist. Novit. 3119:1–86.

12. Johanson Z, Manzanares E, Underwood C, Clark B, Fernandez V, Smith M (2020) Evolution of the Dentition in Holocephalans (Chondrichthyes) Through Tissue Disparity. Integrative and Comparative Biology, 60(3), 630–643. 10.1093/icb/icaa093

13. Smith M, Underwood C, Goral T, Healy C, Johanson Z (2019) Growth and mineralogy in dental plates of the holocephalan Harriotta raleighana (Chondrichthyes): Novel dentine and conserved patterning combine to create a unique chondrichthyan dentition. Zoological Letters, 5(1), 11. 10.1186/s40851-019-0125-3

14. Zhu Y, Li Q, Lu J, Chen Y, Wang J, Gai Z, Zhao W, Wei G, Yu Y, Ahlberg PE, Zhu M (2022) The oldest complete jawed vertebrates from the early Silurian of China. Nature, 609(7929), 954–958. 10.1038/s41586-022-05136-8

15. Stahl, BJ (1999). Handbook of Paleoichthyology Vol. IV. Chondrichthyes III. Holocephali

16. Finarelli JA, Coates MI (2011) First tooth-set outside the jaws in a vertebrate. Proc R Soc B 279:775–779. 10.1098/rspb.2011.1107

17. Grogan ED (1993) The structure of the holocephalan head and the relationships of the Chondrichthyes [PhD dissertation]. (Duquesne University, Pittsburgh, PA).

18. Grogan ED, Lund R, Greenfest-Allen E (2012) The origin and relationships of early chondrichthyans. In Carrier JC, Musick JA, Heithaus MR, eds. Biology of Sharks and Their Relatives, Second Edition. (CRC Press, Boca Raton, FL).

19. Berio F, Debiais-Thibaud M (2021) Evolutionary developmental genetics of teeth and odontodes in jawed vertebrates: A perspective from the study of elasmobranchs. J Fish Biol 98:906–918. 10.1111/jfb.14225

20. Fraser GJ, Hulsey CD, Bloomquist RF, Uyesugi K, Manley NR, Streelman JT (2009) An ancient gene network is co-opted for teeth on old and new jaws. PLoS Biol 7:e1000031. 10.1371/journal.pbio.1000031

21. Haridy Y, Gee BM, Witzmann F, Bevitt JJ, Reisz RR (2019) Retention of fish-like odontode overgrowth in Permian tetrapod dentition supports outside-in theory of tooth origins. Biol Lett 15:20190514. 10.1098/rsbl.2019.0514

22. Nicklin EF, Cohen KE, Cooper RL, Mitchell G, Fraser GJ (2024) Evolution, development, and regeneration of tooth-like epithelial appendages in sharks. Dev Biol 516:221–236. 10.1016/j.ydbio.2024.08.009

23. Smith MM, Coates MI (2000) Evolutionary origins of teeth and jaws: Developmental models and phylogenetic patterns. In Teaford MF, Meredith Smith M, Ferguson MWJ, eds. Development, Function, and Evolution of Teeth, pp. 133–151 (Cambridge University Press, Cambridge, UK). 10.1017/CBO9780511542626.010

24. Fraser GJ, Hulsey CD (2020) Biology at the Cusp: Teeth as a Model Phenotype for Integrating Developmental Genomics, Biomechanics, and Ecology. Integrative and Comparative Biology, 60(3), 559–562. 10.1093/icb/icaa104

25. Hulsey CD, Cohen KE, Johanson Z, Karagic N, Meyer A, Miller CT, Sadier A, Summers AP, Fraser GJ (2020) Grand challenges in comparative tooth biology. Integr Comp Biol 60:563–580. 10.1093/icb/icaa038

26. Rasch LJ, Martin KJ, Cooper RL, Metscher BD, Underwood CJ, Fraser GJ (2016) An ancient dental gene set governs development and continuous regeneration of teeth in sharks. Dev Biol 415:347–370. 10.1016/j.ydbio.2016.01.038

27. Tucker AS, Fraser GJ (2014) Evolution and developmental diversity of tooth regeneration. Semin Cell Dev Biol 25–26:71–80. 10.1016/j.semcdb.2013.12.013

28. Brownstein CD, Near TJ, Dearden RP (2024) The Palaeozoic assembly of the holocephalan body plan far preceded post-Cretaceous radiations into the ocean depths. Proceedings of the Royal Society B: Biological Sciences, 291(2033), 20241824. 10.1098/rspb.2024.1824

29. Venkatesh B, Lee AP, Ravi V, Maurya AK, Lian MM, Swann JB, Ohta Y, Flajnik MF, Sutoh Y, Kasahara M, Hoon S, Gangu V, Roy SW, Irimia M, Korzh V, Kondrychyn I, Lim ZW, Tay BH, Tohari S, … Warren WC (2014) Elephant shark genome provides unique insights into gnathostome evolution. Nature, 505(7482), 174–179. 10.1038/nature12826

30. Fraser GJ, Hamed SS, Martin KJ, Hunter KD (2019) Shark tooth regeneration reveals common stem cell characters in both human rested lamina and ameloblastoma. Scientific Reports, 9(1), 15956. 10.1038/s41598-019-52406-z

31. Huysseune A (2006) Formation of a successional dental lamina in the zebrafish (Danio rerio): Support for a local control of replacement tooth initiation. The International Journal of Developmental Biology, 50(7), 637–643. 10.1387/ijdb.052098ah

32. Smith MM, Fraser GJ, Mitsiadis TA (2009) Dental lamina as source of odontogenic stem cells: Evolutionary origins and developmental control of tooth generation in gnathostomes. Journal of Experimental Zoology Part B: Molecular and Developmental Evolution, *312B*(4), 260–280. 10.1002/jez.b.21272

33. Cooper RL, Nicklin EF, Rasch LJ, Fraser GJ (2022) Teeth outside the mouth: The evolution and development of shark denticles. bioRxiv. 10.1101/2022.07.13.499989

34. Schnetz L, Pfaff C, Kriwet J (2016) Tooth development and histology patterns in lamniform sharks (Elasmobranchii, Lamniformes) revisited. J Morphol 277:1584–1598. 10.1002/jmor.20597

35. Reif WE (1980) A mechanism for tooth pattern reversal in sharks: The polarity switch model. Wilhelm Roux’s Archives of Developmental Biology, 188(2), 115–122. 10.1007/BF00848802

36. Reif WE (1982) Evolution of Dermal Skeleton and Dentition in Vertebrates: The Odontode Regulation Theory. In M. K. Hecht, B. Wallace, & G. T. Prance (Eds.), Evolutionary Biology (pp. 287–368). Springer US. 10.1007/978-1-4615-6968-8_7

37. Cooper RL, Thiery AP, Fletcher AG, Delbarre DJ, Rasch LJ, Fraser GJ (2018) An ancient Turing-like patterning mechanism regulates skin denticle development in sharks. Sci Adv 4:eaau5484. 10.1126/sciadv.aau5484

38. Coates, MI, Tietjen K, Johanson Z, Friedman M, Sang S. (2021) The cranium of Helodus simplex (Agassiz, 1838) revised. In Janvier P, Denton JSS (Eds). Ancient Fishes and their Living Relatives. A Tribute to John G. Maisey (Pp 193-204).

39. De Beer GR, Moy-Thomas JA, Goodrich ES (1937) V I.—On the skull of Holocephali. Philosophical Transactions of the Royal Society of London. Series B, Biological Sciences, 224(514), 287–312. 10.1098/rstb.1935.0001

40. Carrier JC, Musick JA, Heithaus MR (Eds.). (2004). Biology of sharks and their relatives. CRC Press.

41. Nakayama N, Matsunuma M, Endo H. (2020) A preliminary review and in situ observations of the spookfish genus *Harriotta* (Holocephali: Rhinochimaeridae). Ichthyological Research 67(1):82–91. 10.1007/s10228-019-00703-y

42. Vandenplas S, De Clercq A, Huysseune A. (2014) Tooth replacement without a dental lamina: The search for epithelial stem cells in *Polypterus senegalus*. Journal of Experimental Zoology Part B: Molecular and Developmental Evolution322(5):281–293. 10.1002/jez.b.22577

43. Fraser GJ, Standing A, Underwood C, Thiery AP. (2020) The dental lamina: An essential structure for perpetual tooth regeneration in sharks. Integrative and Comparative Biology 60(3):644–655. 10.1093/icb/icaa102

44. Leydig F. (1851) Zur Anatomie und Histologie der *Chimaera monstrosa*. *Archiv für Anatomie,* Physiologie und Wissenschaftliche Medizin 1851:258–268.

45. Lieberman DE, Hall BK. (2006) The evolutionary developmental biology of tinkering: An introduction to the challenge. In: Tinkering: The Microevolution of Development, pp. 1–19 (John Wiley & Sons, Ltd, Hoboken, NJ). 10.1002/9780470319390.ch1

46. Kikinis R, Pieper SD, Vosburgh KG. (2014) 3D Slicer: A platform for subject-specific image analysis, visualization, and clinical support. In: Jolesz FA (Ed). Intraoperative Imaging and Image-Guided Therapy, pp. 277–289 (Springer, New York, NY).

47. Rolfe S, Davis C, Maga AM. (2021) Comparing semi-landmarking approaches for analyzing three-dimensional cranial morphology. American Journal of Physical Anthropology 175(1):227–237. 10.1002/ajpa.24214

