## Supplementary figures and images for "Teeth Outside the Jaw: Evolution and Development of the Toothed Head Clasper in Chimaeras"

### Supplemental Figure 1

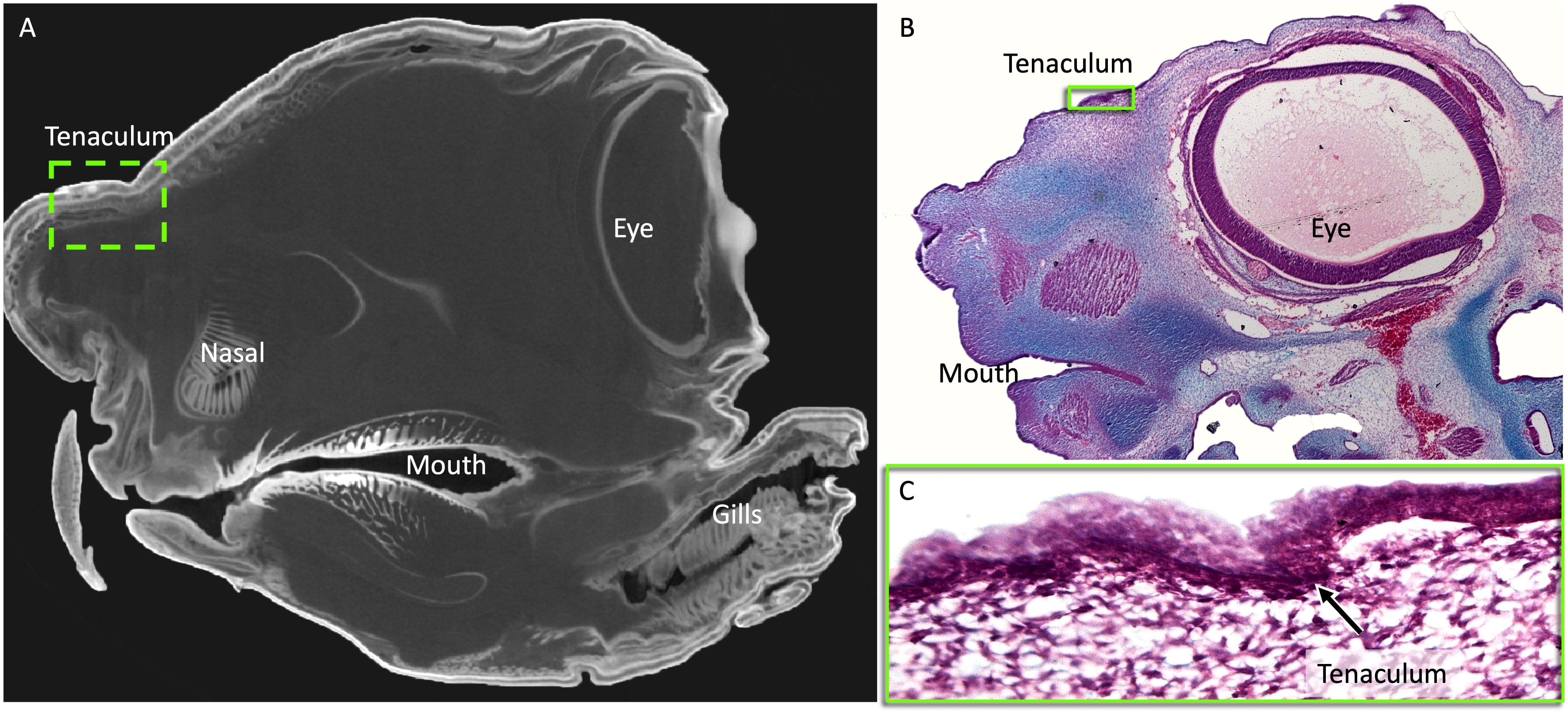

### Supplemental Figure 2

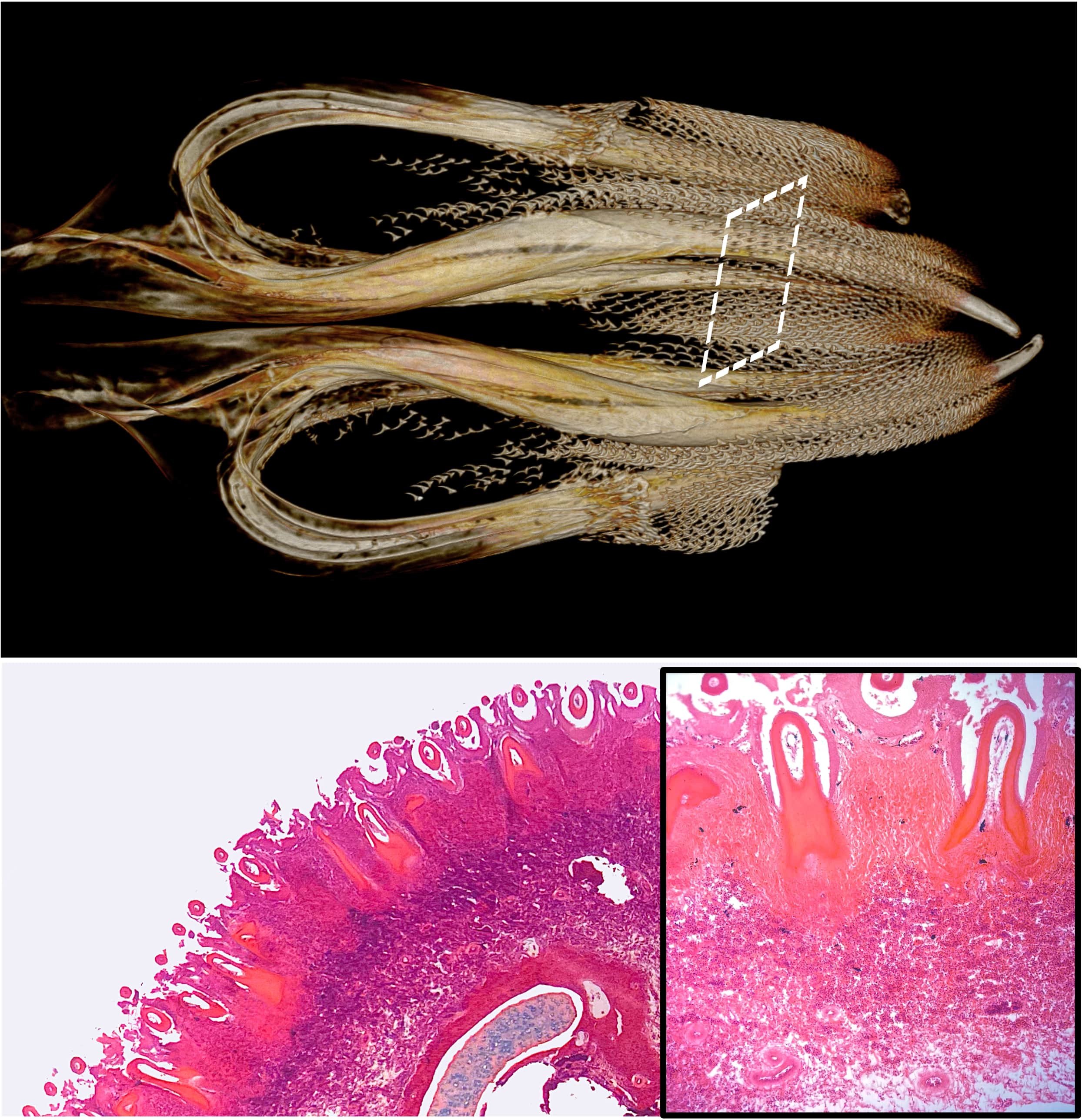
